# Bipedalism in Mexican Albian lizard (Squamata) and the locomotion type in other Cretaceous lizards

**DOI:** 10.1101/2021.04.01.437947

**Authors:** Damián Villaseñor-Amador, Nut Xanat Suárez, J. Alberto Cruz

## Abstract

Representative locomotion types in lizards include terrestrial, arboreal, grass swimmer, sand swimmer and bipedal. Few studies explain the locomotion habit of extinct lizards, and even less asses those of bipedal ones. Here, we use quantitative methods to infer the type of locomotion of two Albian Mexican lizards (Lower Cretaceous) and three Cretaceous lizards from Brazil, North America and Spain, assessing the similarities of the hindlimb-forelimb length ratio amongst extinct and extant species. Additionally, an ancestral character state reconstruction analysis was performed, to evaluate the evolution of lizard locomotion habits. The species *Huehuecuetzpalli mixtecus* was bipedal while *Tijubina pontei* was facultative bipedal, *Hoyalacerta sanzi, Tepexisaurus tepexii* and *Polyglyphanodon sternbergi* cannot be differentiated amongst terrestrial or arboreal with the approach used in this work. The ancestral character state reconstruction analysis showed a terrestrial ancestral locomotion type, with a basal character state of hindlimbs longer than forelimbs. Equal length between hind and forelimbs appear to be a derivate state that evolved multiple times in lizard evolutionary history.

## 1. INTRODUCTION

Lizard bipedalism is found in nine of the 46 extant known families, such as Iguania (Agamidae, Crotaphytidae, Iguanidae, Liolaemidae, Phrynosomatidae, Tropiduridae), Lacertoidea (Lacertidae, Teiidae) and Anguimorpha (Varanidae) (Clemente, 2014; Pyron, 2017; Vitt and Caldwell, 2014). The emergence of bipedalism is believed to have been an exaptation, as a trait to acquire greater maneuverability as speed increased (Clemente, 2014). However, succeeding lineages exploited this trait because of its evolutionary advantages, suggesting a possible adaptative radiation in some groups (Clemente, 2014). Recent works point to the emergence of bidepalism in Squamata in the Lower Cretaceous somewhere during the Aptian-Albian (Simoes et al., 2015; Lee et al., 2018). The most ancient inferred bipedal lizard, due to its hindlimb-forelimb ratio, is *Tijubina pontei* from the Cretaceous of Brazil (Aptian-Albian ∼113 Ma), a species closely related to Polyglyphanodontidae (Simões et al., 2015), an extinct family basal to Lacertoidea (Pyron et al., 2017). Direct fossil evidence of bipedalism in lizards was reported by Lee et al. (2018), who found ichnofossils of lizard footprints belonging to *Sauripes hadongensis*, from 110 Ma ago in the Cretaceous of South Korea.

*Huehuecuetzpalli mixtecus* is a fossil lizard from 105 Ma ago of the Cretaceous Albian of Mexico (Reynoso, 1998). The mention of it being a bipedal lizard exists (Reynoso, 1998; Reynoso and Cruz, 2014), but there is not any study supporting this statement. *H. mixtecus* has been placed at the base of the Iguania clade (Pyron, 2017), alongside *Hoyalacerta* from the Lower Cretaceous of Spain (Evans and Barbadillo, 1999), becoming the sister group to all iguanids (Pyron et al. 2017). *Huehuecuetzpalli mixtecus* was found in the Tlayúa quarry, Tepexi de Rodríguez, Puebla (Reynoso, 1998), a fossiliferous site regarded as Lagerstätte for its excellent fossil preservation (Espinosa-Arrubarrena and Applegate, 1996). The Tlayúa quarry is a small outcrop located in the west side of Tepexi de Rodriguez municipality, Puebla, México, at 18°35’7.24” N and 97°54’38.09” W, 1740 masl, with an age about 105 Ma (Benammi et al., 2006). The fossil lepidosaurians of Tlayúa are represented by two rhynchocephalian sphenodotids (*Ankylosphenodon pachyostosus* and *Pamizinsaurus tlayuensis*) and the lizards *Huehuecuetzpalli mixtecus* and *Tepexisaurus tepexii* (Reynoso and Cruz, 2014) (Figure 1 A, B). *Tepexisaurus tepexii* had short limbs and tail, indicative of crawling movements (Reynoso and Callison, 2000), and *H. mixtecus* had long hindlimbs and tail, which has led to believe it had a bipedal locomotion, with tail autotomy (Reynoso, 1998; Reynoso and Cruz, 2014). Nevertheless, the locomotion inferences of these lizards have only been made using uniformitarianism (Reynoso and Cruz, 2014), without carrying out a quantitative analysis. Using uniformitarianism and quantitative data we can explain Earth history in terms of gradual change by processes observed today (Polly and Spang, 2002), which includes evolutive inferences derived from paleontological records. In this study we quantitatively infer the locomotion type of *H. mixtecus* and *T. tepexii*, and the significance of these within the evolution of bipedalism in Squamata by comparing them to other extinct and extant lizards.

**Figure 1.**
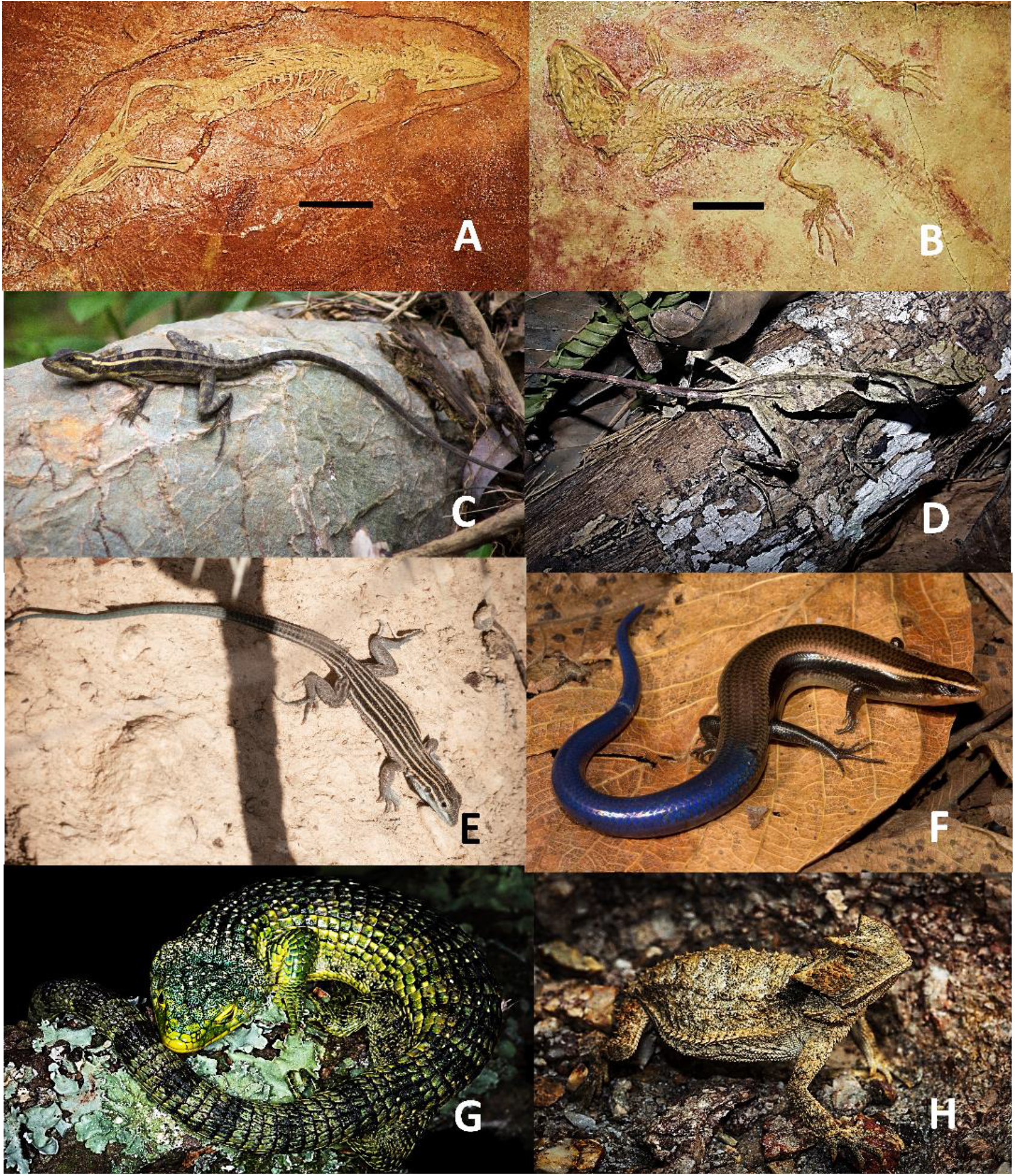
Cretaceous lizards from the Tlayúa Quarry, Mexico (A, B), and locomotion types of extant lizards. Bipedal (C, D), sand swimmer (E), grass swimmer (F), arboreal (G) and terrestrial (H). A) *Huehuecuetzpalli_mixtecus* (IGM 7389), B) *Tepexisaurus tepexii* (IGM 7466), C) *Basiliscus vittatus*, D) *Corythopanes hernandezi*, E) *Aspidoscelis inornata*, F) *Plestiodon lotus*, G) *Abronia graminea* and H) *Phrynosoma sherbrookei*. Photographs by Eric Centenero Alcalá. Scale bar = 3 cm.

## 2. MATERIAL AND METHODS

Photographs were taken of the forelimbs (humerus, radius, ulna) and hindlimbs (femur, tibia, fibula) of 36 existing species of lizards, all with their respective measurement scale. The specimens belong to the osteological reference collection of the paleontology laboratory of the FCB, BUAP (BUAPALO, Table S1). For the digital capture, we used a VE-LX1800 camera mounted in a Miotic SMZ-168 stereoscopic microscope. Bone length was measured with the program Image J v1.8.0 (Ferreira and Rasband, 2012). The work by Gans et al. (2008) was used for identification of disarticulated lizard limb bones. Our current sample of organisms is not very extensive; however recent osteological studies with small sample sizes due to the availability of collection specimens have been known to yield good results with fewer than 30 specimens (Yildirim et al., 2017; Ledesma and Scarpetta, 2018; Paparella et al., 2020).

The locomotion habit by which extant species were catalogued included: bipedal, arboreal, terrestrial, sand swimmer and grass swimmer (Figure 1). Bipedalism involves moving on the hindlimbs alone (Russell and Bels, 2001); many lizards are known to run bipedally, but unlike theropod dinosaurs or birds they do not have bipedal striding gaits (Aerts et al., 2003). The arboreal term is used to describe tree-dweller lizards (Fischer et al., 2010), while ground-dwellers are defined as terrestrial (Russell and Bels 2001). If a ground-dweller lizard shows serpentine movements it will be called sand swimmer if it dwells on sandy environments, or grass swimmer if it is found on grasslands (Maladen, 2009).

Regarding extinct species, holotype high resolution images were consulted for *H. mixtecus* (IGM 7389, 4185) and *T. tepexii* (IGM 7466), both belonging to the paleontological collection of the Institute of Geology, UNAM (Figure 1A, B). The limb bone length was obtained following the same procedure than before. *Polyglyphanodon sternbergi, Hoyalacerta sanzi* and *Tijubina pontei* limb lengths were taken from Gilmore (1942), Evans and Barbadillo (1999) and Bonfim-Júnior and Rocha-Barbosa (2006), respectively.

The hindlimb-forelimb length ratio was calculated as the sum of the femur and tibia longitudes (hindlimb length) divided by the sum of the humerus and ulna longitudes (forelimb length), in mm. Values equal to one, indicated limbs of equal length. Values higher than one showed longer hindlimbs, while values below one showed longer forelimbs. A simple linear regression analysis was performed to evaluate hindlimb-forelimb length correlation. Other measures used were the femur/humerus, tibia/ulna, and fibula/radius ratios which together with the hindlimb-forelimb length ratio were used to construct a Principal Component Analysis (PCA). All analyses were carried out in Rstudio (Rstudio Team 2020; http://www.rstudio.com/).

To analyze the evolution of bipedalism, we used the squamate phylogeny of Pyron et al. (2017). Serpentes and Amphisbaenia groups were excluded, in order to work only with lizards. This phylogeny was chosen because it includes *Huehuecuetzpalli, Hoyalacerta* and *Polyglyphanodon. Tijubina pontei* was placed alongside *Polyglyphanodon*, according to the classification by Simões et al. (2015). A pruned 40 taxa ultrametric tree was obtained. For each taxon, the corresponding species name, hindlimb-forelimb length ratio and locomotion type were provided (Table S1). It is known that analyses based on osteological data of extinct and extant organisms tend to have relatively small sample sizes (Cardini and Elton, 2007; Duan et al., 2020) because of rarity and collection difficulty (Brown and Vavrek, 2015). In our case we have a limited number of specimens due to availability constrains (see above). However scarce fossil taxa may be, it can improve the accuracy of phylogenetic analysis of morphological datasets (Koch et al., 2020). Two ancestral character state reconstructions were carried out. The first one was issued discrete values: the types of locomotion for each of the extant and fossil lizards. The extinct species locomotion type was implied from the statistical analysis of this work (Table 1). To identify the likely plesiomorphic condition for the type of locomotion, of the studied lizard taxa, a stochastic mapping for discrete phylogeny traits was performed (Bollback, 2006; Revell, 2012), with a bootstrapping of 1000 iterations. The analysis was carried out under the maximum likelihood approach (Bollback, 2006) and an equal-rates model (Pagel, 1994; Pagel, 1999) was selected as the model of character state evolution, with equal probability for any change. The equal-rates model recognizes *a priori* information, such as the fossil records used in this work (Pagel, 1994; Pagel, 1999; Skinner, 2010).

**Table 1.**
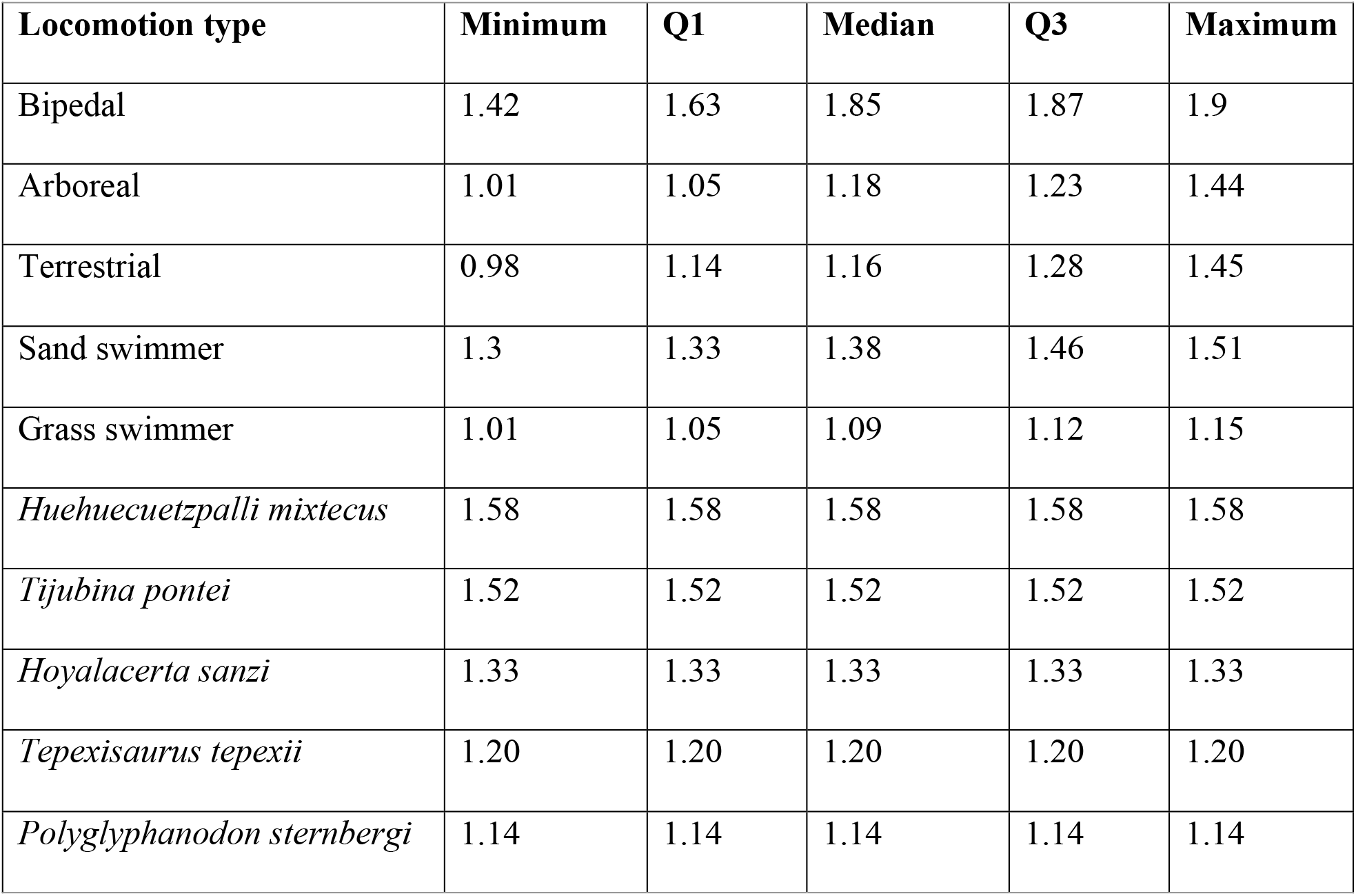
Minimum, maximum, median and quartiles (Q), in millimeters, of the relationship between hind (femur + tibia) and forelimb (humerus + ulna) bone length.

For the second ancestral character state reconstructions analysis, a continuous trait was given: the hindlimb-forelimb length ratio (Table 1). Classification of species as bipedal, or other locomotion habit, was mapped onto a phylogenetic tree. In order to characterize hindlimb-forelimb length ratio evolution, seven models of evolution were tested, and the one providing the best fit was chosen. The evolution models are listed as follows: *i*) a single-rate Brownian motion model (BM1); *ii*) an Ornstein–Uhlenbeck model with one optimum for all lizard species (OU1); *iii*) a Brownian motion model that allowed for separate rates for each locomotion habit regime (BMS); *iv*) an OU_M_ model that allows for different optima for each regime. And three OU models allowing for *v*) different Brownian motion rates (OU_MV_), *vi*) different strength of selection parameters (OU_MA_) or *vii*) different Brownian motion rates and different strength of selection parameters (OU_MVA_). The evolution model that best described continuous traits evolving under discrete selective pressures (O’Meara et al., 2006), was chosen as the one providing the best Akaike Information Criterion corrected for small sample sizes (AICc) fit (O’Meara et al., 2006). In order to infer the historic change of lizard hindlimb-forelimb proportion, a phylogenetic mapping estimating the transitional states, using maximum likelihood approach (Revell, 2012) and Felsenstein methodology (Felsenstein, 1985), was carried out.

All analyses were performed in R v4.0.3 (R Core Team, 2020). For stochastic and transitional states phylogenetic mapping, we used the *phytools* R package v0.6.99 (Revell, 2012) and for the evolution model testing, the *OUwie*.*R* function from the *OUwie* R package v2.6 (Beaulieu et al., 2012; Beaulieu and O’Meara, 2019; Beaulieu and O’Meara, 2021). and for the delta AICc scores the *akaike*.*weights* function from the *qpcR* R package v1.4-1 (Spiess, 2018).

## 3. RESULTS

### 3.1 Extant and extinct lizard locomotion habits

Hindlimb-forelimb length ratio shows that grass swimmer species are the lizards with the most similar limb length (Q1 – Q3 = 1.05 – 1.12). Of all the groups, terrestrial (Q1 – Q3 = 1.14 – 1.28) and arboreal (Q1 – Q3 = 1.05 – 1.23) lizards display the largest length variation.

Sand swimmer species appear to have slightly longer hindlimbs (Q1 – Q3 = 1.33 – 1.46), while bipedal ones show the highest ratio values of them all (Q1 – Q3 = 1.63 – 1.87) (Table 1, Figure 2A). For the Cretaceous lizards, it is not possible to determine the locomotion habit of *Tepexisaurus tepexii* and *Polyglyphanodon sternbergi*, due to its hindlimb-forelimb length ratio (1.19 and 1.14, respectively) being inside the terrestrial-arboreal interval (Figure 2A).

**Figure 2.**
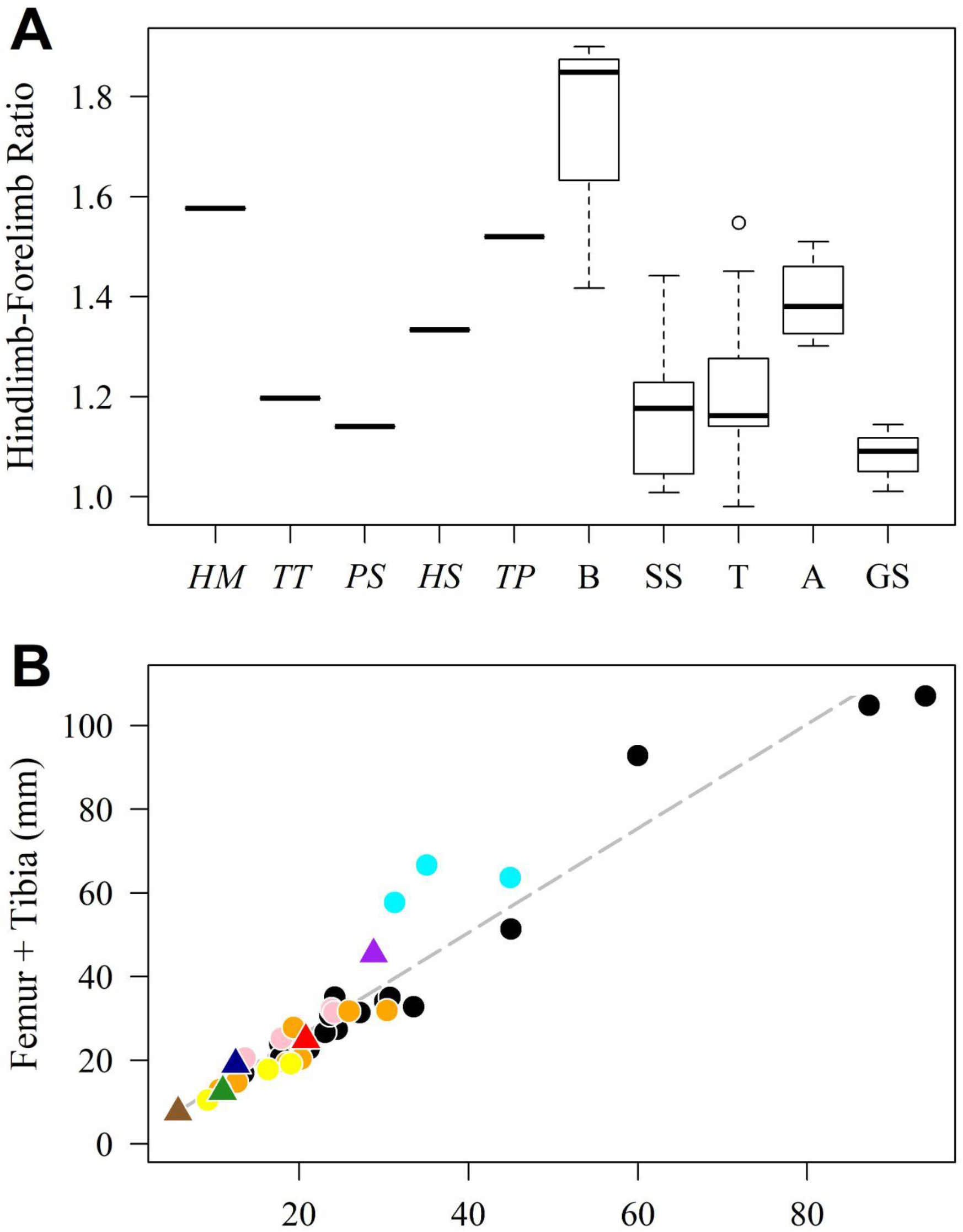
A) Hindlimb-forelimb length ratio boxplot, for the Cretaceous lizards *Huehuecuetzpalli mixtecus* (*HM*), *Tepexisaurus tepexii* (*TT*), *Polyglyphanodon sternbergi* (*PS*), *Hoyalacerta sanzi* (*HS*), *Tijubina pontei* (*TP*) and the locomotion types of 36 extant species. Abbreviations: B stands for bipedal; SS = sand swimmer; T = terrestrial; A = arboreal; GS = grass swimmer. B) Linear regression for the longitude of hindlimb (femur + tibia) by forelimb (humerus + ulna) of extant (circles) and fossil (triangles) lizards. Black stands for terrestrial locomotion habit; sky blue = bipedal; pink = sand swimmer; orange = arboreal; yellow = grass swimmer; purple = *Huehuecuetzpalli mixtecus*; red = *Tepexisaurus tepexii*; dark blue = *Tijubina pontei*; green = *Polyglyphanodon sternbergi*; gray = *Hoyalacerta sanzi*.

Nevertheless, a terrestrial locomotion type is assumed for both species, according to what has been published (Gilmore, 1942; Reynoso and Callison, 2000). *Hoyalacerta sanzi* has marginally longer hindlimbs (1.33), which fall in between the quartiles of the sand swimmer, terrestrial and arboreal locomotion habit (Figure 2A). Both *Tijubina pontei* and *Huehuecuetzpalli mixtecus* hindlimb-forelimb length ratios (Table 1) are above sand swimmer variation values (Min – Max = 1.3 – 1.51) but are encompassed by bipedal ones (Min -Max = 1.42 – 1.9).

Simple linear regression analysis shows a positive correlation between hind and forelimbs (R^2^=0.55, *P*<0.0001). All extant lizards are near or below the regression line, except the bipedal species and one terrestrial species (Figure 2B). These last four have longer hindlimbs than forelimbs. Of the Cretaceous fossils, *Huehuecuetzpalli mixtecus* raises above the regression line, grouped with the extant bipedal lizards. Meanwhile, *Tijubina pontei* barely goes above the line, near the sand swimmer locomotion type lizards. *Hoyalacerta sanzi, Polyglyphanodon sternbergi* and *Tepexisaurus tepexii* are inside the regression line, being unable with this analysis to determine their locomotion habit.

PCA shows to *Polyglyphanodon sternbergi, Tepexisaurus tepexi*, and *Hoyalacerta sanzi* inside the terrestrial and arboreal variation, *Huehuecuetzpalli mixtecus* is clear inside the bidpedal locomotion type and *Tijubina pontei* is outside of all locomotion type (Figure 3A). This led us to infer that *Huehuecuetzpalli mixtecus* was a bipedal lizard supported by all analyses, like extant lizards *Basilicus, Laemanctus*, and *Corytophanes* (Figure 3B). *Tijubina pontei* was possibly a facultative bipedal lizard because the hindlimb-forelimb length ratios (Table 1) are above sand swimmer values close to bipedal variation values (Figure 2A); however, it is possible that bidepalism is not supported by the regression or PCA analyzes (Figure 2, 3). We could not infer the locomotion type for *Polyglyphanodon sternbergi, Tepexisaurus tepexii*, and *Hoyalacerta sanzi*, since the fossils fell within terrestrial and arboreal locomotion (Figure 2, 3).

**Figure 3.**
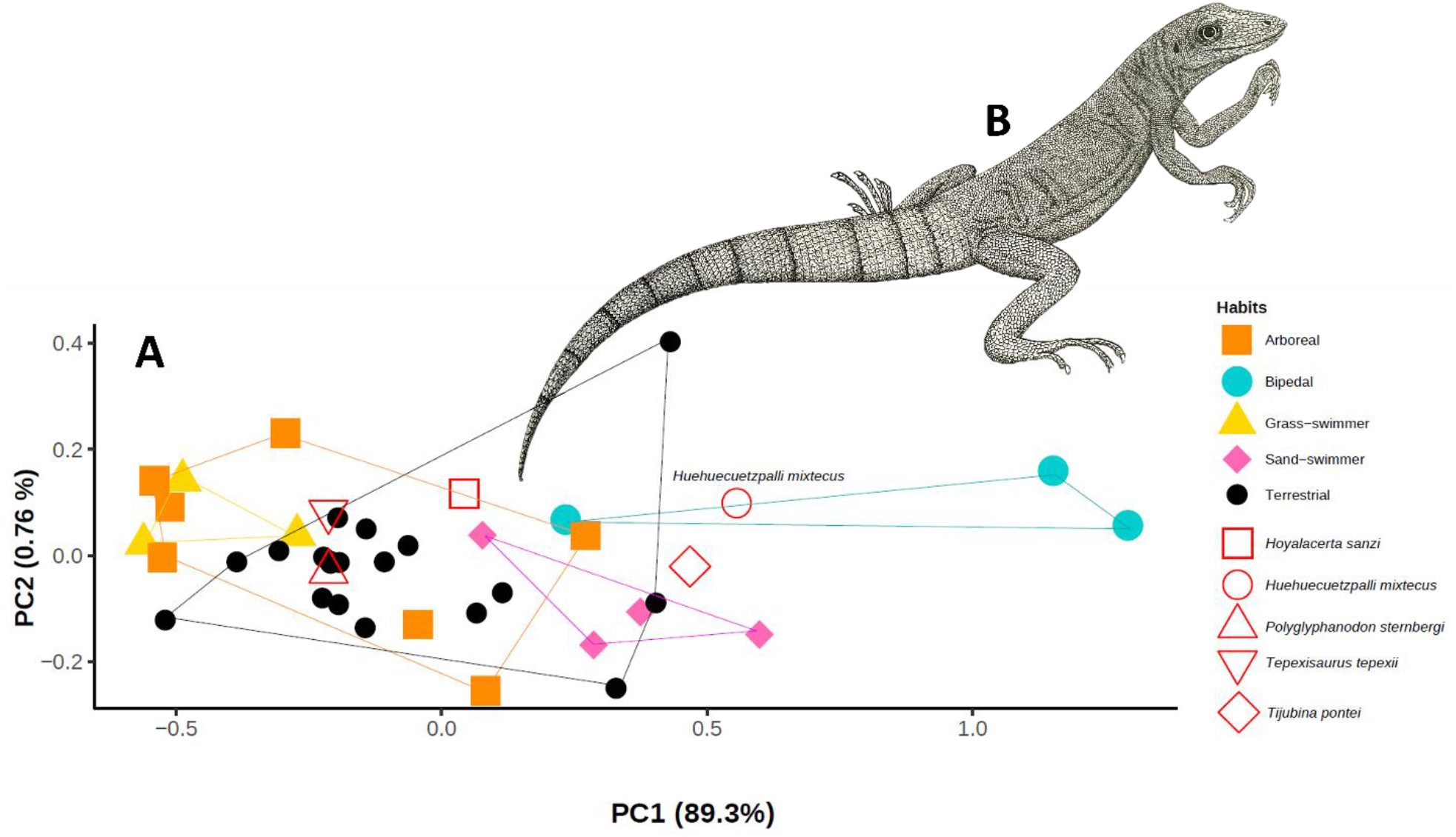
A) PCA of the locomotion types in extant (filled figures) and extinct (empty figures) lizards. B) Reconstruction of a lizard *Huehuecuetzpalli mixtecus* in bipedal position, iguanid body and varanid head, based on the description of Reynoso and Cruz (2014). Drawn by Gerardo García Demeneghi.

### 3.2 Evolution of lizard bipedalism

Ancestral state reconstruction of discrete character infers terrestrial locomotion habit as the most likely condition at the base of the phylogeny (51% inference value, Figure 3B). Other habits show similar inference values (bipedal = 12%, arboreal = 10%, sand swimmer = 17% and grass swimmer = 10%). No group displays a unique locomotion type, for all the habits emerge rapidly in different groups (Figure 4B), and terrestrial locomotion habit seems to be the ancestral condition at internal nodes of the phylogeny. Nonetheless, bipedalism only develops in Lacertoidea with *Tijubina pontei*, and Iguania with *Huehuecuetzpalli mixtecus, Basiliscus vittatus, Corytophanes hernandezi* and *Laemanctus serratus*.

**Figure 4.**
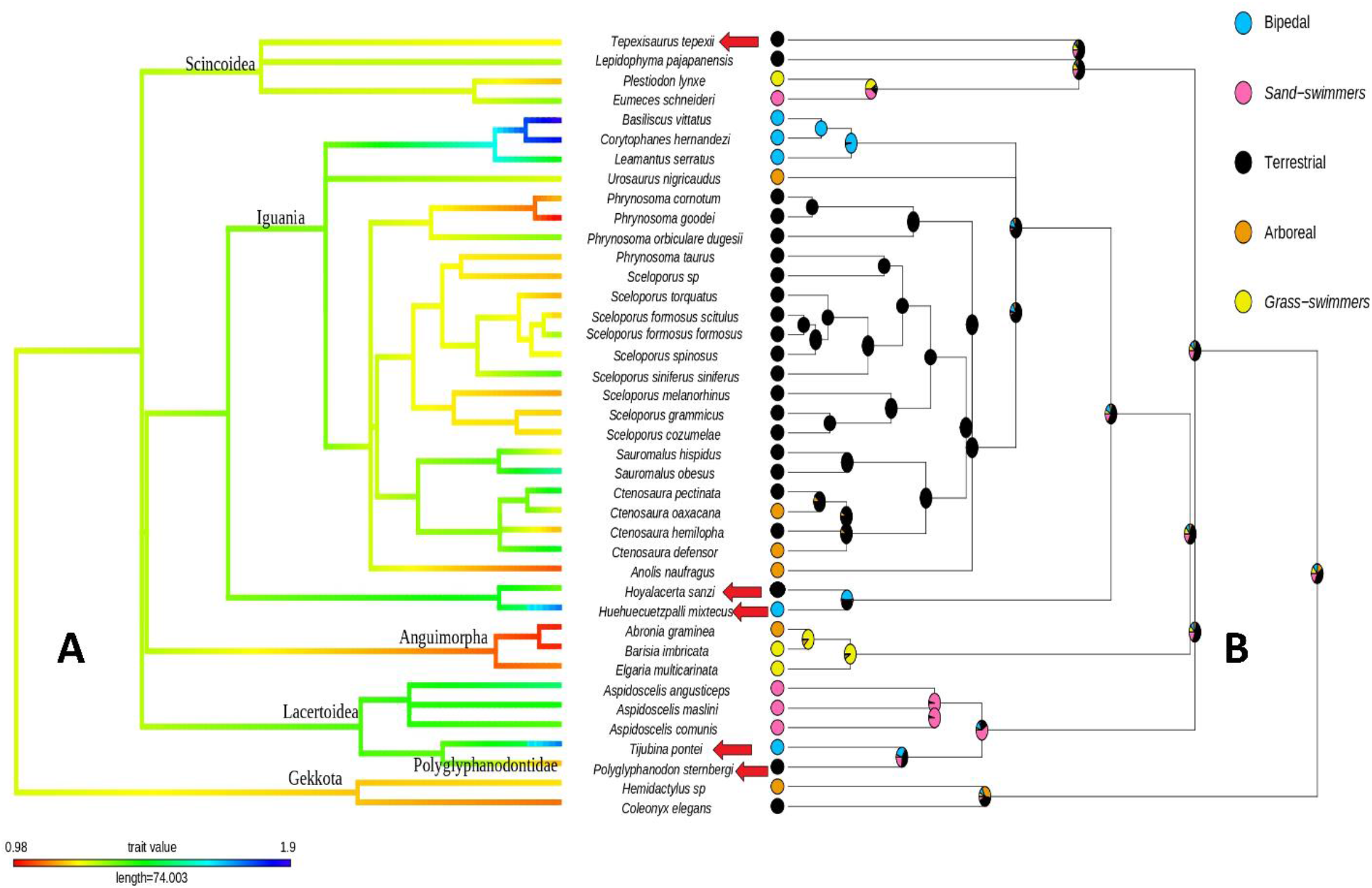
Phylogenetic tree and evolution model of lizard hindlimb-forelimb length ratio. A) Values close to one (orange - red) show a similar length ratio for arboreal (e.g. *Abronia gramminea*), grass swimmer (e.g. *Barisia imbricata*) and terrestrial (e.g. *Phrynosoma goodei*) species. Values different from one (light-dark blue) indicate longer hindlimbs for bipedal (e.g. *Basiliscus vittatus, Huehuecuetzpalli mixtecus, Tijubina pontei*), sand swimmer (*Aspidoscelis angusticeps*) and terrestrial (e.g. *Sauromalus obesus*) lizards. Intermediate values (green - yellow) denote marginally longer forelimbs for terrestrial and sand swimmer locomotion habits. B) Mapping of the locomotion types of extant and fossil (arrows) lizard species. The ancestral state is inferred to be terrestrial at the base of the phylogeny (51%) in relation to other habits: bipedal (12%), arboreal (10%), sand swimmer (17%) and grass swimmer (10%).

The evolution model that provided the best fit for characterizing the hind-forelimb length relationship, was the OU_MA_(Table 2). Which indicates that strength of selection (α) and the optima (θ) differ between the selective regimes (locomotion habits), but not so the rate of stochastic motion (σ^2^). The terrestrial state had the highest strength of selection (0.5408), followed by bipedal (0.0135), grass-swimmer (0.0047), arboreal (0.0005) and sand-swimmer (0.0002). The optima for terrestrial lizards was 1.14, for all others it was 0. Longer hindlimbs are only present in modern day family Corytophanidae (*Basiliscus, Corytophanes* and *Laemanctus*) and the fossil lizards *Huehuecuetzpalli mixtecus* (Iguania) and *Tijubina pontei* (Lacertoidea). Species with slightly longer hindlimbs were the extant terrestrial iguanas *Sauromalus obesus* and *Ctenosaura pectinata*, as well as the sand swimmer teiid *Aspidoscelis angusticeps* (Figure 3A). Limb length relationship points to a basal character state of marginally longer hindlimbs. Equal length between hind and forelimbs appear to be a derivate state that evolved multiple times in lizard evolutionary history, among terrestrial (e.g. *Phrynosoma goodei*), arboreal (e.g. *Abronia gramminea*) and sand swimmer (e.g. *Barisia imbricata*) species (Figure 3A). On the other hand, the development of larger hindlimbs can be discerned in bipedal extant (*Basiliscus, Corytophanes* and *Laemanctus*) and extinct (*H. mixtecus* and *T. pontei*) taxa, as well as some sand swimmer (*Aspidoscelis angusticeps*) and terrestrial (*Sauromalus obesus*) species (Figure 3A).

**Table 2.** Akaike Information Criterion corrected for small sample sizes (AICc) and Bayesian Information Criteria (BIC) for the seven evolutionary models tested in this work. OU_MA_ model best describes the evolution of hindlimb-forelimb length relationship for extant and extinct lizards. Likelihood, AICc and BIC scores were determined using the function *OUwie* in the *OUwie* R package (Beaulieu & O’Meara, 2021). Delta AICc scores were then calculated using the function *akaike*.*weights* in the *qpcR* R package (Spiess, 2018).

| Model | -lnL | AICc | ΔAICc | BIC |
| --- | --- | --- | --- | --- |
| BM1 | 8.65 | -13.29 | -12.97 | -9.91 |
| BMS | 13.43 | -12.32 | 703.95 | -4.73 |
| OU1 | 13.06 | -19.45 | 696.82 | -15.05 |
| OU<sub>M</sub> | 25.21 | -32.93 | 683.34 | -24.60 |
| OU<sub>MV</sub> | 28.56 | -25.69 | 690.58 | -16.54 |
| OU<sub>MA</sub> | **373.85** | **-716.27** | **0** | **-707.12** |
| OU<sub>MVA</sub> | 27.99 | -5.97 | 710.30 | -0.64 |

## 4. DISCUSSION

The hindlimb-forelimb length ratio indicates a clear difference between lizard bipedalism and other locomotion types (Figures 2, 3). Differences were also shown among sand swimmer and grass swimmer species, but were not between terrestrial and arboreal ones, due to their wide variation range (Table 1, Figure 2, 3). It was inferred that *Huehuecuetzpalli mixtecus* exhibited a bipedal locomotion, while *Tijubina pontei* exhibited a facultative bipedal locomotion type, due to the hindlimb-forelimb length ratio values not being as high as those of extant lizards (Table 1, Figures 2, 3). Because length values of *Hoyalacerta sanzi, Tepexisaurus tepexii* and *Polyglyphanodon sternbergi* are similar between hind and forelimbs, it was not possible to discern if these species were either terrestrial or arboreal (Table 1, Figure 2, 3). In order to distinguish between terrestrial and arboreal locomotion it will be necessary to carry out further studies, either with claw (Herrel et al., 2002; Zani, 2008) or hip (Fischer et al., 2010; Russell and Bels, 2001) measurements.

Linear bone limb measurement, as well as hindlimb-forelimb length ratio, have already been used to infer bipedal locomotion habits, such as those in the extinct reptiles *Eudibamus cursoris* from the Permian of Germany (Berman et al., 2000) and *Langobardisaurus* from the Triassic of Italy (Renesto et al., 2002). Multiple times, these parameters have also been used to infer extant lizard locomotion types (Abou Egla et al., 2008; Druelle et al., 2019; Foster et al., 2018; Herrel et al., 2002; Snyder, 1962), demonstrating their effectivity at studying the locomotion of extinct organisms.

Lizard bipedalism is the easiest locomotion type to differentiate, due to the fact that hindlimbs will always be longer than forelimbs (Grinham and Norman, 2020; Schuett et al., 2009; Snyder, 1962). This allowed us to infer bipedalism and the facultative bipedalism of the Cretaceous lizards *Huehuecuetzpalli mixtecus* and *Tijubina pontei*, respectively. It is facultative because their hindlimb-forelimb length ratio is less than extant bipedal lizards, such as *Basilicus, Corytophanes* and *Leamanctus* (Figure 2, 3). Bipedal locomotion of *T. pontei* was possibly a consequence of morphology (hindlimb longer than forelimbs) and acceleration, allowing more maneuverability as speed increased (Clemente, 2014).

With the inclusion in the analysis of other Cretaceous lizard fossils (*Hoyalacerta sanzi, Huehuecuetzpalli mixtecus, Polyglyphanodon sternbergi, Tepexisaurus tepexii* and *Tijubina pontei*), terrestrial locomotion habit was inferred to be the ancestral trait. This is in accordance with the hypothesis that several locomotion habits are derivates from the quadrupedal terrestrial type (Russell and Bels, 2001). Strength of selection values (α) of the OU_MA_ model further support this: the highest α value was for the terrestrial locomotion habit (0.5408). Bipedal locomotion had the second highest a value (0.0135). The hindlimb-forelimb ratio optimum (θ) was the terrestrial habit (1.14) while all the others locomotion types tended to 0. It is known that OU_MA_, OU_MV_ and OU_MVA_ models tend to overestimate α and underestimate θ, but this happens even with big sample sizes (i.e. 512 taxa) (Beaulieu et al., 2012). Nevertheless, our interpretations should be considered with caution.

*Huehuecuetzpalli mixtecus* and *T. pontei* are basal to Iguania and Lacertoidea which are two out of the three clades that possess modern day bipedal lizards (the third being Anguimorpha) (Clemente, 2014). This strengthens the hypothesis of lizard bipedalism emergence as an exaptation, which later favored certain organisms as an evolutive advantage (Clemente, 2014). Among these are some corytophanids, agamids and varanids (Clemente, 2014; Schuett et al., 2009). We acknowledge that most of the specimens used in our study correspond to specific lineages in Iguania and this overrepresentation could lead to biased conclusions. Nonetheless the objective of our study was to evaluate the significance of *H. mixtecus* and *T. tepexii* in the evolution of bipedalism in lizards, because *H. mixtecus* is basal to Iguania (Pyron et al., 2017). Ancestral character state reconstructions of the hindlimb-forelimb length ratio showed that the ancestral locomotion type belonged to a terrestrial lizard with marginally longer hindlimbs (Figure 3A). This is consistent with the oldest fossil record of lizard-like diapsid limbs, such as those of the species: *Paleagama vielhaueri* and *Saurosternon bainii* from the Permian-Jurassic (Carroll, 1975) and *Hongshanxi xiei* from the Jurassic (Dong et al., 2019); all three species had longer hindlimbs (Table 3). From the ancestral state of hindlimbs slightly longer than forelimbs, the extant lizard limb proportions arose. For example, those of the species with same limb lengths (arboreal, terrestrial and grass swimmer), and those of the taxa with longer hindlimbs (bipedal, sand swimmer and, occasionally, terrestrial). However, further research is needed to explain the origin and diversification of lizard locomotion habits.

**Table 3.** Hind and forelimb length of the Permian-Jurassic diapsids *Paleagama vielhaueri, Saurosternon bainii* and *Hongshanxi xiei*. Note the higher hindlimb values.

| **Fossil** | **Forelimb**<br>**(humerus + ulna length)** | **Hindlimb**<br>**(femur + tibia length)** |
| --- | --- | --- |
| *Paleagama vielhaueri* | 38.2 mm | 55.4 mm |
| *Saurosternon bainii* | 28.8 mm | 38.8 mm |
| *Hongshanxi xiei* | 13.8 mm | 22.5 mm |

## 5. CONCLUSIONS

Bipedal locomotion habit is inferred for the Cretaceous lizard *Huehuecuetzpalli mixtecus* and facultative bipedal locomotion habit is inferred for *Tijubina pontei*.

The locomotion type of *Hoyalacerta sanzi, Tepexisaurus tepexii* and *Polyglyphanodon sternbergi* cannot be differentiated between terrestrial or arboreal, with the approach used in this work. We infer that the last common lizard ancestor had a terrestrial locomotion habit, with hindlimbs slightly longer than forelimbs, whereas hindlimbs equally elongated as forelimbs or strongly longer than forelimbs is a derived condition.

Lizards with longer hindlimbs can be interpreted as being bipedal or facultative bipedal. By contrast, grass swimmers have the same length between their limbs. Sand swimmers have long hindlimbs, but without being particularly long as in the case of bipedal lizards. Finally, the most variation in hindlimb-forelimb length ratio was observed for the terrestrial and arboreal habits. However, because of the overrepresentation of Iguanid lineages in our study and specimen availability limitations, further research with more diverse samples is needed.

## ACKNOWLEDGEMENTS

We are thankful to the Program “Haciendo Ciencia en la BUAP Primavera XIV 2019” from the Vicerrectoría de Investigación y Estudios de Posgrado, Benemérita Universidad Autónoma de Puebla (BUAP), for the support given to undergraduates DVA (No.DGDC/VIEP/0141/2019) and NXS (No.DGDC/VIEP/0142/2019). We are grateful to Eric Centenero Alcalá for supplying the photographs of extant lizards, Gerardo García Demeneghi for drawing the *Huehuecuetzpalli mixtecus* reconstruction and two anonymous peer reviewers, for their invaluable remarks who enriched the current paper.

## SUPPLEMENTARY MATERIAL

Table S1. Excel spreadsheet with bones measurements, type of locomotion and catalog number of extinct and extant lizards used in this study.

## Notes

### Competing Interest Statement

The authors have declared no competing interest.

