## Supplementary material for "Bipedalism in Mexican Albian lizard (Squamata) and the locomotion type in other Cretaceous lizards": Table S1

Table S1. Measurements, type of locomotion and catalog number of extinct and extant lizards used in this study

| locomotion type | species | catalogue | femur | tibia | fibula | humerus | radius | ulna | fe+t | h+u | fe+t/h+u | fe/h | t/u | fi/r |
|---|---|---|---|---|---|---|---|---|---|---|---|---|---|---|
| Bipedal | *Basiliscus vittatus* | BUAPALO 411 | 36.64 | 30.02 | 30.01 | 19.64 | 14.57 | 15.44 | 66.66 | 35.08 | 1.9002 | 1.8655 | 1.944 | 2.0597 |
| Bipedal | *Corytophanes hernandezi* | BUAPALO 248 | 31.46 | 26.32 | 26.45 | 16.59 | 13.37 | 14.67 | 57.78 | 31.26 | 1.8483 | 1.8963 | 1.7941 | 1.9783 |
| Bipedal | *Leamantus serratus* | BUAPALO 172 | 35.08 | 28.59 | 27.79 | 24.84 | 19.8 | 20.1 | 63.67 | 44.94 | 1.4167 | 1.4122 | 1.4223 | 1.4035 |
| Grass-swimmer | *Barisia imbricata* | BUAPALO 173 | 11.75 | 7.49 | 7.56 | 11.09 | 7.13 | 7.95 | 19.24 | 19.04 | 1.0105 | 1.0595 | 0.9421 | 1.0603 |
| Grass-swimmer | *Elgaria multicarinata* | BUAPALO 247 | 10.93 | 6.88 | 6.74 | 9.35 | 6.8 | 6.98 | 17.81 | 16.33 | 1.0906 | 1.1689 | 0.9856 | 0.9911 |
| Grass-swimmer | *Plestiodon lynxe* | BUAPALO 210 | 6.39 | 4.14 | 4.04 | 5.32 | 3.25 | 3.88 | 10.53 | 9.2 | 1.1445 | 1.2011 | 1.0670 | 1.2430 |
| Sand-swimmer | *Aspidoscelis angusticeps* | BUAPALO 250 | 11.37 | 9.18 | 9.22 | 8.01 | 5.19 | 5.6 | 20.55 | 13.61 | 1.5099 | 1.4194 | 1.6392 | 1.7764 |
| Sand-swimmer | *Aspidoscelis comunis* | BUAPALO 164 | 17.48 | 14.71 | 15.52 | 13.66 | 9.42 | 10.17 | 32.19 | 23.83 | 1.3508 | 1.2796 | 1.4464 | 1.6475 |
| Sand-swimmer | *Aspidoscelis maslini* | BUAPALO 251_196 | 13.73 | 11.54 | 11.78 | 10.07 | 7.06 | 7.85 | 25.27 | 17.92 | 1.4101 | 1.3634 | 1.4700 | 1.6685 |
| Sand-swimmer | *Eumeces schneideri* | BUAPALO 407 | 18.96 | 12.45 | 13.71 | 13.98 | 9.31 | 10.16 | 31.41 | 24.14 | 1.3011 | 1.3562 | 1.2253 | 1.4726 |
| Arboreal | *Abronia graminea* | BUAPALO 303 | 12.3 | 8.06 | 8.28 | 11.54 | 7.28 | 8.65 | 20.36 | 20.19 | 1.0084 | 1.0658 | 0.9317 | 1.1373 |
| Arboreal | *Anolis naufragus* | BUAPALO 375 | 18.68 | 13.22 | 12.85 | 16.48 | 12.26 | 13.88 | 31.9 | 30.36 | 1.0507 | 1.1334 | 0.9524 | 1.0481 |
| Arboreal | *Ctenosaura defensor* | BUAPALO 200 | 15.59 | 12.3 | 12 | 11.22 | 8.78 | 8.12 | 27.89 | 19.34 | 1.4420 | 1.3894 | 1.5147 | 1.3667 |

| | | | | | | | | | | | | | | |
|---|---|---|---|---|---|---|---|---|---|---|---|---|---|---|
| Arboreal | *Ctenosaura oaxacana* | BUAPALO 191 | 17.08 | 14.73 | 14.9 | 14.91 | 11.09 | 10.98 | 31.81 | 25.89 | 1.2286 | 1.1455 | 1.3415 | 1.3435 |
| Arboreal | *Hemidactylus sp* | BUAPALO 194 | 9.46 | 5.4 | 6.29 | 7.03 | 5.38 | 5.6 | 14.86 | 12.63 | 1.1765 | 1.3456 | 0.9642 | 1.1691 |
| Arboreal | *Pachydactylus briboni* | BUAPALO 204 | 11.76 | 7.58 | 7.91 | 10.08 | 7.6 | 8.5 | 19.34 | 18.58 | 1.0409 | 1.1666 | 0.8917 | 1.0407 |
| Arboreal | *Urosaurus nigricaudus* | BUAPALO 287 | 7.13 | 5.9 | 6.1 | 6.32 | 3.89 | 4.29 | 13.03 | 10.61 | 1.2280 | 1.1281 | 1.3752 | 1.5681 |
| Terrestrial | *Coleonyx elegans* | BUAPALO 181 | 12.19 | 10.59 | 10.72 | 10.95 | 9 | 10.22 | 22.78 | 21.17 | 1.0760 | 1.1132 | 1.0362 | 1.1911 |
| Terrestrial | *Sceloporus torquatus* | BUAPALO 175 | 19.06 | 15.05 | 14.6 | 16.85 | 11.02 | 13.25 | 34.11 | 30.1 | 1.1332 | 1.1311 | 1.1358 | 1.3248 |
| Terrestrial | *Sceloporus spinosus* | BUAPALO 195 | 13.89 | 10.5 | 10.5 | 10.94 | 7.93 | 9.35 | 24.39 | 20.29 | 1.2020 | 1.2696 | 1.1229 | 1.3240 |
| Terrestrial | *Sceloporus sp* | BUAPALO 182 | 19.77 | 15.3 | 16.41 | 16.44 | 12.05 | 14.28 | 35.07 | 30.72 | 1.1416 | 1.2025 | 1.0714 | 1.3618 |
| Terrestrial | *Sceloporus siniferus siniferus* | BUAPALO 249 | 12.65 | 11.33 | 11.67 | 10.28 | 6.93 | 7.45 | 23.98 | 17.73 | 1.3525 | 1.2305 | 1.5208 | 1.6839 |
| Terrestrial | *Sceloporus melanorhinus* | BUAPALO 167 | 14.86 | 12.59 | 12.67 | 12.8 | 10.4 | 11.68 | 27.45 | 24.48 | 1.1213 | 1.1609 | 1.0779 | 1.2182 |
| Terrestrial | *Sceloporus grammicus* | BUAPALO 219 | 11.71 | 8.97 | 9.12 | 9.76 | 6.87 | 8.04 | 20.68 | 17.8 | 1.1617 | 1.1997 | 1.1156 | 1.3275 |
| Terrestrial | *Sceloporus formosus scitulus* | BUAPALO 162 | 15.01 | 11.68 | 12.14 | 12.6 | 9.32 | 10.45 | 26.69 | 23.05 | 1.1579 | 1.1912 | 1.1177 | 1.3025 |
| Terrestrial | *Sceloporus formosus formosus* | BUAPALO 165 | 18.3 | 14.17 | 13.58 | 15.17 | 9.87 | 10.04 | 32.47 | 25.21 | 1.2879 | 1.2063 | 1.4113 | 1.3758 |
| Terrestrial | *Sceloporus cozumelae* | BUAPALO 183 | 8.55 | 5.89 | 5.97 | 6.76 | 4.61 | 5.47 | 14.44 | 12.23 | 1.1807 | 1.2647 | 1.0767 | 1.2950 |
| Terrestrial | *Phrynosoma taurus* | BUAPALO 8 | 17.4 | 14.03 | 13.82 | 15.51 | 9.93 | 11.66 | 31.43 | 27.17 | 1.1567 | 1.1218 | 1.2032 | 1.3917 |

| | | | | | | | | | | | | |
|---|---|---|---|---|---|---|---|---|---|---|---|---|
| Terrestrial | *Phrynosoma orbicular edugesii* | BUAPALO 246 | 17.31 | 13.47 | 13.23 | 13.57 | 8.94 | 10.05 | 30.78 | 23.62 | 1.3031 | 1.2756 | 1.3402 | 1.4798 |
| Terrestrial | *Phrynosoma goodei* | BUAPALO 7 | 18.63 | 14.21 | 14.97 | 19.09 | 12.62 | 14.42 | 32.84 | 33.51 | 0.9800 | 0.9759 | 0.9854 | 1.1862 |
| Terrestrial | *Lepidophyma pajapanensis* | BUAPALO 166 | 10.02 | 6.96 | 7.38 | 8.03 | 5.9 | 5.4 | 16.98 | 13.43 | 1.2643 | 1.2478 | 1.2888 | 1.2508 |
| Terrestrial | *Sauromalus obesus* | BUAPALO 202 | 54.35 | 38.5 | 38.76 | 30.29 | 25.88 | 29.7 | 92.85 | 59.99 | 1.5477 | 1.7943 | 1.2962 | 1.4976 |
| Terrestrial | *Ctenosaura hemilopha* | BUAPALO 344 | 63.02 | 44.09 | 46.67 | 55.33 | 34.16 | 38.64 | 107.11 | 93.97 | 1.1398 | 1.1389 | 1.1410 | 1.3662 |
| Terrestrial | *Ctenosaura pectinata* | BUAPALO 197 | 19.45 | 15.66 | 15.52 | 14.23 | 9.87 | 9.97 | 35.11 | 24.2 | 1.4508 | 1.3668 | 1.5707 | 1.5724 |
| Terrestrial | *Phrynosoma cornotum* | BUAPALO 6 | 29.74 | 21.66 | 22.77 | 24.72 | 17 | 20.28 | 51.4 | 45 | 1.1422 | 1.2030 | 1.0680 | 1.3394 |
| Terrestrial | *Sauromalus hispidus* | BUAPALO 11 | 62.18 | 42.71 | 43.27 | 50.22 | 31.26 | 37.08 | 104.89 | 87.3 | 1.2014 | 1.2381 | 1.1518 | 1.3841 |
| Fossil | *Tepexisaurus tepexii* | IGM 7466 | 14.2 | 10.7 | 9.9 | 11.4 | 8.3 | 9.4 | 24.9 | 20.8 | 1.1971 | 1.2456 | 1.1382 | 1.1927 |
| Fossil | *Huehuecuetzpalli mixtecus* | IGM 7389 | 24.7 | 20.7 | 20.3 | 15.7 | 12.9 | 13.1 | 45.4 | 28.8 | 1.5763 | 1.5732 | 1.5801 | 1.5736 |
| Fossil | *Polyglyphanodon sternbergi* | USNM 15477 | 7.23 | 5.29 | 5.6 | 6.1 | 4.13 | 4.88 | 12.52 | 10.98 | 1.1402 | 1.1852 | 1.0840 | 1.3559 |
| Fossil | *Hoyalacerta sanzi* | LH 11000 | 4.4 | 3.2 | 2.6 | 3.2 | 2 | 2.5 | 7.6 | 5.7 | 1.3333 | 1.375 | 1.28 | 1.3 |
| Fossil | *Tijubina pontei* | MPSC-V 010 | 10 | 9 | 8 | 7 | 5.3 | 5.5 | 19 | 12.5 | 1.52 | 1.4285 | 1.6363 | 1.5094 |
